# Auxin-triggered changes in the Arabidopsis root tip (phospho)proteome reveal novel root growth regulators

**DOI:** 10.1101/2021.03.04.433936

**Authors:** Natalia Nikonorova, Evan Murphy, Cassio Flavio Fonseca de Lima, Shanshuo Zhu, Brigitte van de Cotte, Lam Dai Vu, Daria Balcerowicz, Lanxin Li, Xiangpei Kong, Gieljan De Rop, Tom Beeckman, Jiří Friml, Kris Vissenberg, Peter C. Morris, Zhaojun Ding, Ive De Smet

## Abstract

Auxin plays a dual role in growth regulation and, depending on the tissue and concentration of the hormone, it can either promote or inhibit division and expansion processes in plants. Recent studies revealed that, beyond transcriptional reprogramming, alternative auxin-controlled mechanisms regulate root growth. Here, we explored the impact of different auxin concentrations on the root tip proteome and phosphoproteome, generating a unique resource. From the phosphoproteome data we pinpointed (novel) growth regulators, such as the RALF34-THE1 module. Our results together with previously published studies suggest that auxin, H^+^-ATPases, cell wall modifications and cell wall sensing receptor-like kinases are tightly embedded in a pathway regulating cell elongation. Furthermore, our study assigned a novel role to MKK2 as a regulator of primary root growth and a (potential) regulator of auxin biosynthesis and signalling, and suggests the importance of the MKK2 Thr^31^ phosphorylation site for growth regulation in the *Arabidopsis* root tip.

**ONE SENTENCE SUMMARY:** An auxin-triggered Arabidopsis root tip (phospho)proteome reveals novel root growth regulators

## INTRODUCTION

One of the exceptional features that distinguishes plants from animals is the ability to continuously grow throughout their life. Constant root growth is essential for plant survival, water and nutrition uptake and adaptation to environmental stresses (Petricka et al., 2012; Satbhai et al., 2015). The source of this ceaseless growth is in the meristems that initiate the formation of new tissues and organs. The root meristem has a well-defined structure with stereotypical patterns of cell types along radial and longitudinal axes (Benková and Hejátko, 2009) and a stem cell niche to maintain growth (Dolan et al., 1993; van den Berg et al., 1998; Stahl et al., 2009; Heidstra and Sabatini, 2014; Kong et al., 2018). Along the longitudinal axis, the primary root is divided in meristematic, elongation and differentiation zones (Verbelen et al., 2006). For meristem maintenance and continuous root growth, the rate of cell differentiation should be equal to the rate of cell division (Sozzani and Iyer-Pascuzzi, 2014; Pacifici et al., 2015). Such balance is under tight control of multiple developmental triggers with a key role assigned to phytohormones (Benková and Hejátko, 2009; Pacifici et al., 2015; Di Mambro et al., 2017).

Phytohormones are naturally occurring molecules that act at very low concentrations as signalling compounds and regulate plant growth and development. One group of hormones named auxins (from the Greek – ‘to grow’) earned its name because of its ability to induce growth responses in plants. Establishment and maintenance of auxin gradients in plants control the two main components of growth – cell division and cell expansion (Rahman et al., 2007; Perrot-Rechenmann, 2010; Wang and Ruan, 2013). Auxin plays a dual role in growth regulation and, depending on the tissue and concentration of the hormone, it can both promote and inhibit division and expansion processes in plants (Rayle et al., 1970; Mulkey et al., 1982; Evans et al., 1994; Barbez et al., 2017). Recent studies showed that increased cellular auxin levels result in dramatically enhanced root cell elongation, altered expression of cell wall remodelling genes and reduced cell wall arabinogalactan complexity (Pacheco-Villalobos et al., 2013; Pacheco-Villalobos et al., 2016). A positive effect of auxin on growth was hypothesized by the acid growth theory (Hager, 2003; Takahashi et al., 2012; Du et al., 2020). This theory postulates that auxin triggers the activation of plasma membrane (PM)-localized H^+^-ATPases (proton pumps), resulting in acidification of the apoplast, activation of cell wall-loosening enzymes, and turgor pressure-mediated cell expansion. However, the acid growth theory in roots still remains the subject of debate. Recent studies showed that higher cellular auxin levels in *Brachypodium* roots were not related to proton pump activation or elevated proton excretion (Pacheco-Villalobos et al., 2016). However, auxin was shown to be important for apoplast acidification and stimulation of cell expansion in *Arabidopsis* root (Barbez et al., 2017). Furthermore, relatively high concentrations of exogenous auxin as well as endogenous elevation of auxin levels lead to root growth reduction (Rahman et al., 2007; Barbez et al., 2017). Notably, both, auxin treatment and activation of auxin biosynthesis result in transient apoplast alkalinisation (Barbez et al., 2017). However, the role of the apoplastic pH in root growth regulation remains unclear since other cellular mechanisms downstream of auxin have been proposed, including microtubule re-arrangements (Chen et al., 2014) or vacuolar fragmentation (Löfke et al., 2015; Scheuring et al., 2016). In addition, the inhibitory effect of high auxin levels on primary root growth could be a result of auxin–ethylene cross-talk as auxin stimulates ethylene biosynthesis (Abel et al., 1995; Woeste et al., 1999) and ethylene leads to cell wall alkalinisation (Staal et al., 2011) and inhibition of root cell elongation (Markakis et al., 2012).

Canonical auxin signalling starts with auxin binding to the receptor complex, followed by modulation of gene transcription and protein abundance (Tan et al., 2007; Chapman and Estelle, 2009; Slade et al., 2017). However, recent studies also showed an alternative mechanism in roots involving intra-cellular auxin perception, but not transcriptional reprogramming (Fendrych et al., 2018). Although our understanding of the effects of auxin on *Arabidopsis* root growth at the protein and phosphorylation level is increasing (Mattei et al., 2013; Zhang et al., 2013; Slade et al., 2017; Cao et al., 2019; Huang et al., 2019; Lv et al., 2020), it still remains incomplete. To address this gap in our knowledge, we explored the impact of different auxin concentrations on the root tip proteome and phosphoproteome.

## RESULTS AND DISCUSSION

### Establishment of optimum auxin levels for repression and promotion of primary root growth

To establish the optimum auxin concentrations affecting *Arabidopsis* primary root growth in our hands, seedlings were grown vertically for 11 days after germination in the absence or continuous presence of 0.1, 10 and 100 nM naphthalene-1-acetic acid (NAA, a synthetic auxin analogue). This root assay revealed that 0.1 nM of NAA promoted primary root growth; however, constant exposure to 10 and 100 nM NAA caused primary root growth reduction (**Supplementary Figure S1A**). Taken together, this root assay indicated that at low concentration auxin promotes primary root growth and that at high concentration auxin caused growth reduction. However, so far, it remained largely unclear what the underlying molecular mechanisms controlling these opposite responses are.

### Proteome and phosphoproteome profiling to unravel concentration-dependent root growth auxin response

To gain insight into molecular changes associated with auxin-triggered growth regulation, we focused on changes in the proteome and phosphoproteome in the *Arabidopsis* root tip. Specifically, *Arabidopsis* seedlings were grown vertically on solid ½ MS medium containing different concentrations of NAA (mock, 0.1 nM, 10 nM and 100 nM) and at 11 days after germination the root tip (1 cm) was harvested in 5 biological repeats. We chose a 1 cm root tip as in this zone we could capture the effect of auxin on both cell division and cell expansion, but at the same time avoid the region with auxin-induced lateral root formation (**Supplementary Figure S1B**). We furthermore chose 11-day-old seedlings as we detected a high level of difference in primary root growth upon auxin treatment at this age. From these root tips (5 biological replicates), proteins were extracted and used for two parallel analyses: (i) the total proteome, enabling us to identify key proteins responding to auxin concentration gradients, and (ii) the phosphoproteome, allowing us to gain insights into the auxin concentration-dependent phosphorylation events.

The proteome analysis of control and auxin-treated samples resulted in the identification of 3193 protein groups (a protein group includes proteins that cannot be unambiguously identified by unique peptides but have only shared peptides) (**Figure 1 and Supplementary Table S1**). The phosphoproteome analysis led to the identification of 6548 phosphorylated sites (belonging to 2196 proteins) (**Figure 1 and Supplementary Table S2**). Statistical analysis of the proteome data determined 127 differentially abundant proteins after auxin treatment (**Figure 1 and Supplementary Table S1**). In addition, five differentially abundant proteins were identified as ‘unique’, because these were not at all detected in at least one of the treatment conditions (**Figure 1 and Supplementary Table S1**). At the same time analysis of the phosphoproteome dataset revealed 443 differentially abundant phosphopeptides that could be mapped on 346 proteins and 59 ‘unique’ (detected in maximum 1 out of 5 replicates of at least one condition) phosphorylated peptides that are derived from 55 proteins (**Figure 1 and Supplementary Table S2**). Interestingly, our data indicated that auxin-mediated growth responses appeared to be more pronounced at the level of phosphorylation of proteins rather than changes in protein abundance (**Figure 2A-B**). Specifically, the proportion of differentially regulated proteins was only 4% from the total identifications, while this was 18% for the proteins with altered phosphorylation. Furthermore, at the level of protein abundance there were very few differential concentration-specific proteins, while this was more pronounced at the level of the phosphoproteins (**Figure 2C**).

**Figure 1.**
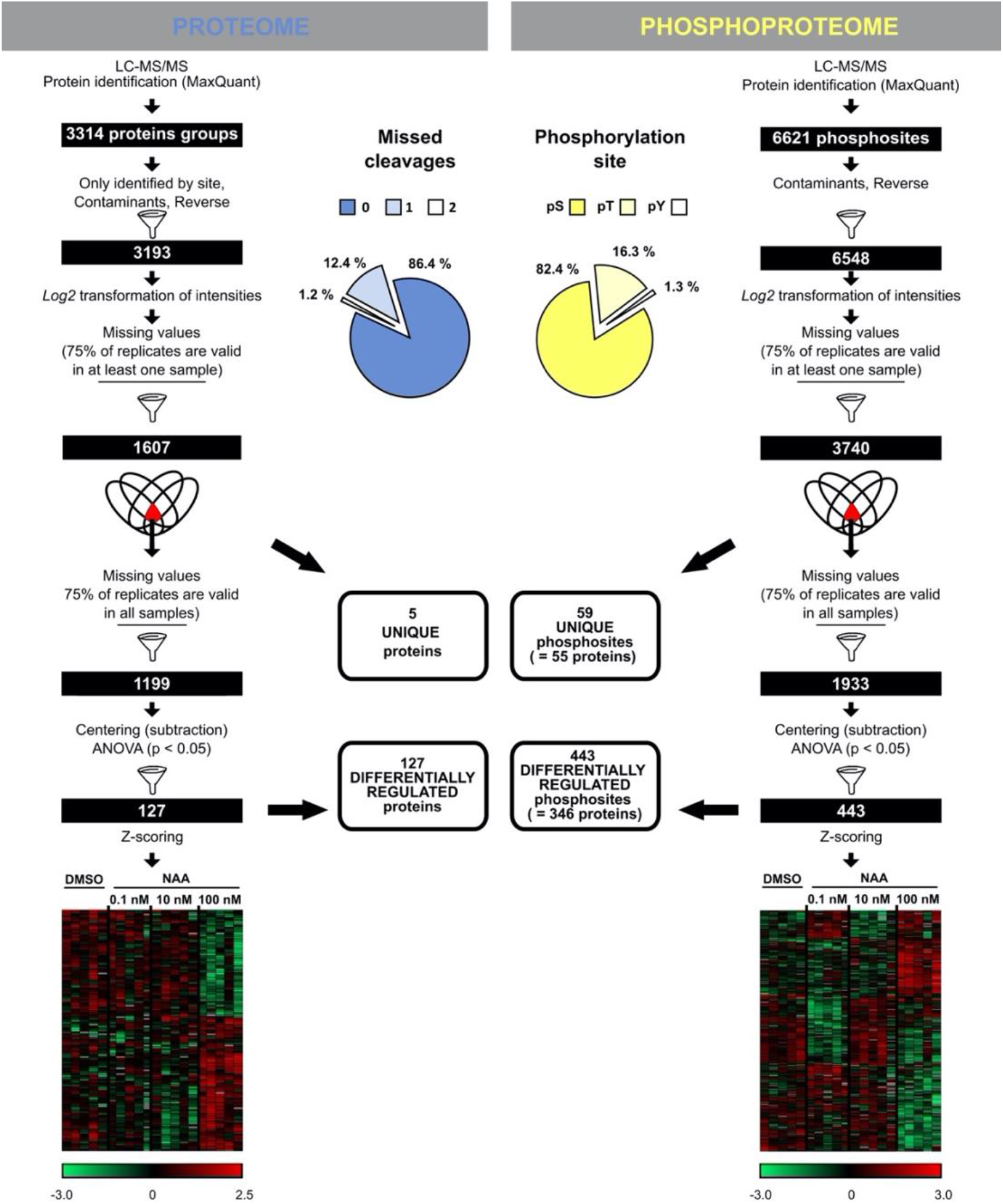
Auxin-triggered protein and phosphoprotein changes. Workflow illustrating the steps to obtain a reliable set of proteins or phospho-sites following LC-MS/MS. Venn diagrams indicate steps where ‘unique’ proteins/phosphosites (with corresponding numbers) were filtered out from the statistical analysis. Heatmaps represent a hierarchical clustering of statistically significant proteins and phosphosites based on Pearson correlation. Centered Z-scored log2-transformed intensity values on heatmaps are color-coded according to provided color gradient scales.

**Figure 2.**
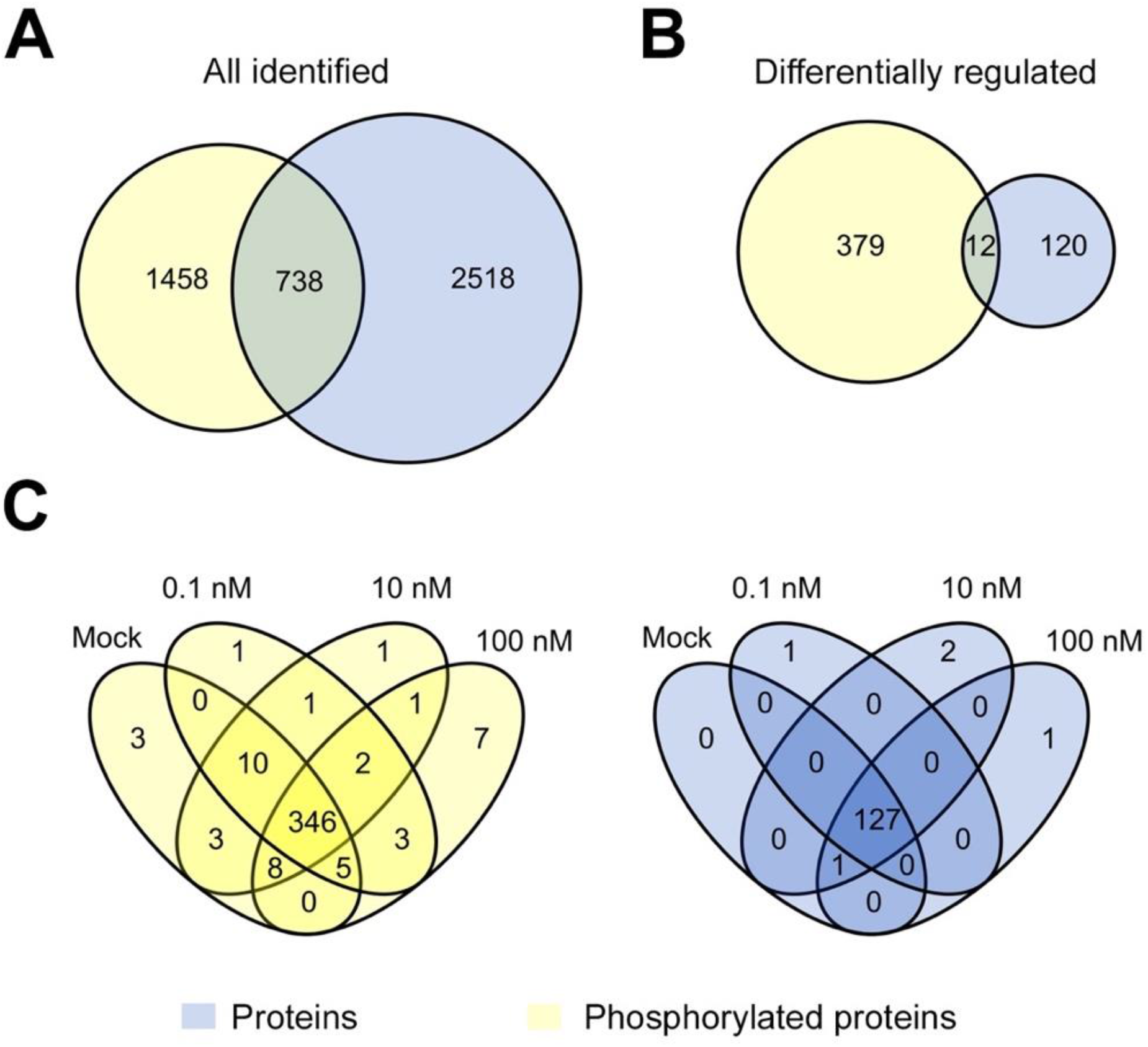
Comparison of auxin proteome and phosphoproteome data. **(A-B)** Overlap of all identified and differentially regulated proteins (blue) and phosphoproteins (yellow) in proteome and phosphoproteome datasets. **(C)** Overlap between differentially regulated (including ‘unique’ ones) proteins (blue) or phosphoproteins (yellow) at different auxin concentrations.

### Auxin-triggered effects on the Arabidopsis root tip proteome

Using the PLAZA 4.0 platform (Van Bel et al., 2018), GO enrichment analysis was conducted on the 132 differentially regulated proteins in the context of a biological process (**Supplementary Figure S2**). The GO enrichment on biological processes showed that differentially regulated proteins were involved in processes such as regulation of cell wall organization and biogenesis, negative regulation of growth, and response to H_2_O_2_ (with a 2-fold log2 enrichment cut-off). Hierarchal clustering of differentially regulated proteins revealed two large clusters of proteins mainly up- or downregulated in samples treated with the growth inhibiting concentration of NAA (100 nM) (**Supplementary Figure S3**). This could explain the enriched GO terms such as negative growth regulation and stress responses.

In this study, we wanted to unravel differences in processes underlying growth promotion and inhibition upon an auxin concentration gradient. Thus, we focused on candidates that were (i) exclusively present/absent in samples treated with 0.1 and 100 nM NAA and (ii) gradually changing their abundance along the increase/decrease of auxin concentration, with both direct and inverse correlation of abundance to the auxin concentration gradient (**Supplementary Figure S4**).

A first group of proteins are potentially involved in growth promoting responses. Two proteins were highly up or down-regulated for the growth-promoting NAA concentration (0.1 nM) and thus potentially involved in positively regulating primary root growth and/or belong to the molecular description of primary root growth. The first one, RAB GTPASE HOMOLOG G3F (RABG3F, AT3G18820) was present only at 0.1 nM NAA (**Supplementary Figure S5A**). Rab GTPases coordinate membrane traffic and function as molecular switches (Stenmark, 2009). RABG3F, a member of the plant Rab7 small GTPase family, localises to prevacuolar compartments and the tonoplast and mediates the final step of vacuolar trafficking. Mutations in RABG3F affect vacuole morphology, inhibited vacuolar trafficking, delayed storage protein degradation and lead to seedling death (Cui et al., 2014). The second one, RNA-BINDING PROTEIN 47B (RBP47B, AT3G19130) was absent only at 0.1 nM NAA (**Supplementary Figure S5B**). RBP47B is a polyadenylate-binding protein involved in RNA binding and aggregation in the cytoplasm and thereby is a part of highly complex and dynamic regulation of mRNA translation, stabilisation, and turnover (Chantarachot and Bailey-Serres, 2018). Also, 32 proteins decreased in abundance along with increasing auxin concentration. This group contained proteins mainly associated with protein folding, regulation of translation, transcription and responses to cytokinin and heat (**Supplementary Figure S6 and Supplementary Table S3**).

A second group of proteins are potentially involved in growth inhibition responses. The only protein that was exclusively present in samples treated with 100 nM NAA was PEROXIDASE 2 (PRX2, AT1G05250), a cationic cell-wall-bound peroxidase involved in lignin biosynthesis (Shigeto et al., 2013; Shigeto et al., 2014) (**Supplementary Figure S5C**). The abundance of 48 proteins was positively correlated with the increasing auxin concentration gradient (direct correlation). This group contained proteins involved in oxidation-reduction and stress responses, carbohydrate metabolism and abscisic acid and gibberellin responses (**Supplementary Figure S6 and Supplementary Table S4**). Among those, several proteins involved in oxidative stress response were identified, including CATALASE 2 (CAT2, AT4G35090), PEROXIDASE 21 (PER21, AT2G37130), PEROXIDASE 57 (PER57, AT5G17820), PEROXIDASE 69 (PER69, AT5G64100) and FE SUPEROXIDE DISMUTASE 1 (FSD1, AT4G25100), which aligns with the fact that the regulation of reactive oxygen species and auxin biology are interconnected (Joo et al., 2001). Peroxidases can play a role in the cross-linking of cell wall hydroxyproline-rich glycoproteins (De Cnodder et al., 2005); but, we also identified other proteins involved in cell wall organisation or biogenesis and carbohydrate metabolism. Altogether our proteome data highlights a tight control of the balance between cell wall loosening and stiffening by auxin in a concentration-dependent manner, and suggests that the regulation of reactive oxygen species seems to play an important role during auxin-mediated primary root growth.

### Auxin-triggered effects on the Arabidopsis root tip phosphoproteome

GO enrichment analysis of biological process terms revealed that proteins with differentially regulated phosphosites under auxin treatment (including ‘unique’ phosphosites) were involved in various processes, including regulation of cell growth (**Supplementary Figure S7**). Hierarchal clustering of 443 differentially abundant phosphosites revealed eight clusters (**Supplementary Figures S8A**). The two biggest clusters contained more than half of the differentially phosphorylated sites (238 sites = 54 %) of which 103 and 135 were specifically up- and downregulated, respectively, in samples treated with the highest auxin concentration. Next, two large clusters representing a third of the differentially abundant phosphosites (135 sites = 31 %) were up- or downregulated at both growth promoting (0.1 nM) and growth inhibiting (100 nM) auxin concentration and likely include general growth regulators not directly linked to growth promotion or inhibition. In addition, two other clusters contained 57 sites that changed their phosphorylation status only after treatment with the growth promoting auxin concentration. Additionally, we clustered differentially abundant phosphosites normalised to protein abundances (**Supplementary Tables S5** and **Supplementary Figure S8B**). The normalisation to protein abundances revealed, for example, differentially phosphorylated phosphopeptides of two 26S proteasome regulatory subunits RPN2a (AT2G32730) and RPN3a (AT1G20200). Phosphosites of both regulatory subunits were up-regulated upon auxin. However, Ser^896^ from RPN2a (AT2G32730) was more phosphorylated at the growth promoting NAA concentration and Ser^14^ from RPN3a - at the growth inhibiting NAA concentration.

A first set of differential phosphosites pinpointed proteins potentially involved in growth promotion responses (**Supplementary Figure S9**). Phosphorylation of SOLUBLE NSF ATTACHMENT PROTEIN RECEPTOR (SNARE) associated Golgi protein family (AT1G71940) at position Ser^17^ was identified only in samples treated with 0.1 nM NAA. Previously, it was shown that auxin controls the abundance of vacuolar SNARE complex components (Löfke et al., 2015). Our data indicates that auxin has an impact on phosphorylation of vacuolar SNAREs by possibly affecting its stability and/or activity. In the proteome data we also detected RABG3F as one of the ‘unique’ proteins for growth promoting responses. Endosomal/vacuolar RAB GTPases and SNARE complexes are functionally connected by tethering complexes to mediate tethering of two membranes before membrane fusion (Wickner and Schekman, 2008). Additionally, ten phosphopeptides were exclusively absent in samples treated only with 0.1 nM NAA (**Supplemental Table S2**). Furthermore, 102 phosphopeptides belonging to 78 phosphoproteins decreased in abundance along with an increasing auxin concentration. This group contained proteins involved in transcription and RNA splicing, and responses to ABA and karrikin; but the function of a big group of phosphoproteins was unknown (**Supplementary Table S6**).

A second set of differential phosphopeptides pinpointed proteins potentially involved in growth inhibition responses (**Supplemental Figure S9**). Nine phosphopeptides were identified only in the sample treated with 100 nM NAA. In addition, 15 phosphopeptides were absent in samples treated with 100 nM NAA. The latter group included a peptide with Ser^218^ from an auxin efflux carrier PIN-FORMED 2 (PIN2, AT5G57090) and a peptide with Ser^159^ from a Major Facilitator Superfamily (MFS) transporter ZINC INDUCED FACILITATOR-LIKE 1 (ZIFL1, AT5G13750). ZIFL1 was shown to indirectly modulate cellular auxin efflux during shootward auxin transport at the root tip, by regulating plasma membrane PIN2 abundance (Remy et al., 2013). Additionally, the abundance of 98 phosphosites derived from 86 phosphoproteins was positively correlated with the auxin concentration gradient (**Supplementary Table S2**). Several phosphoproteins were involved in oxidation-reduction process and stress responses, regulation of transcription and cell cycle, protein phosphorylation and diverse transport-related processes (**Supplementary Table S7**).

### Validation of selected candidates

To validate our proteome and phosphoproteome data sets, we selected a few proteins that showed auxin-mediated differential abundance or phosphorylation and evaluated the growth of respective mutants on auxin.

First, we identified 17 (potential) protein kinases for which phosphorylation sites were significantly regulated by auxin, including four members of MAPK cascades and several receptor (-like) kinases (**Supplementary Table S8**). With respect to the receptor-like kinases, we first focused on THESEUS1 (THE1, AT5G54380), which plays an important role in hypocotyl elongation (Hématy et al., 2007). *THE1* is transcriptionally induced by brassinosteroids (BR), another growth regulating plant hormone (Guo et al., 2009), and *THE1* is expressed in the elongation zone of primary roots (Hématy et al., 2007). The THE1 phosphopeptide containg Ser^668^ was exclusively detected upon auxin treatment and absent in mock samples (**Figure 3A**). The Ser^668^ is located in the protein kinase domain and highly conserved among all the members of the *Catharanthus roseus*-like receptor like kinases 1 (*Cr*RLK1) (**Supplementary Figure S10**). For our analysis, we selected a knockout allele *the1-1* containing a point mutation at a conserved residue of the extracellular domain (Hématy et al., 2007). The primary root length in 11-day-old seedlings of *the1-1* is not significantly longer in control conditions, but is more responsive or less sensitive to the growth promoting or growth repressing NAA treatment, respectively, compared to Col-0 (**Figure 3B**). It should, however, be noted that the difference in length is only 7.1 %, 4.3 % and 9.4 % for 0.1, 10 and 100 nM NAA, respectively. This could possibly be explained by functional redundancy of THE1 with related receptor-like kinases. In this context, we also identified a phosphopeptide belonging to ERULUS (ERU) (**Supplementary Figure S11A**), another member of the CrRLK1 family (Schoenaers et al., 2018). However, based on the *ERU* expression pattern and previously reported phenotypes, the role of ERU is likely only associated with root hair and pollen tube tip-growth (Schoenaers et al., 2017; Schoenaers et al., 2018). Indeed, an *eru* mutant did not display an obvious primary root length phenotype (**Supplementary Figure S11B**), so probably other redundant root-expressed *CrRLK1* family members play a role.

**Figure 3.**
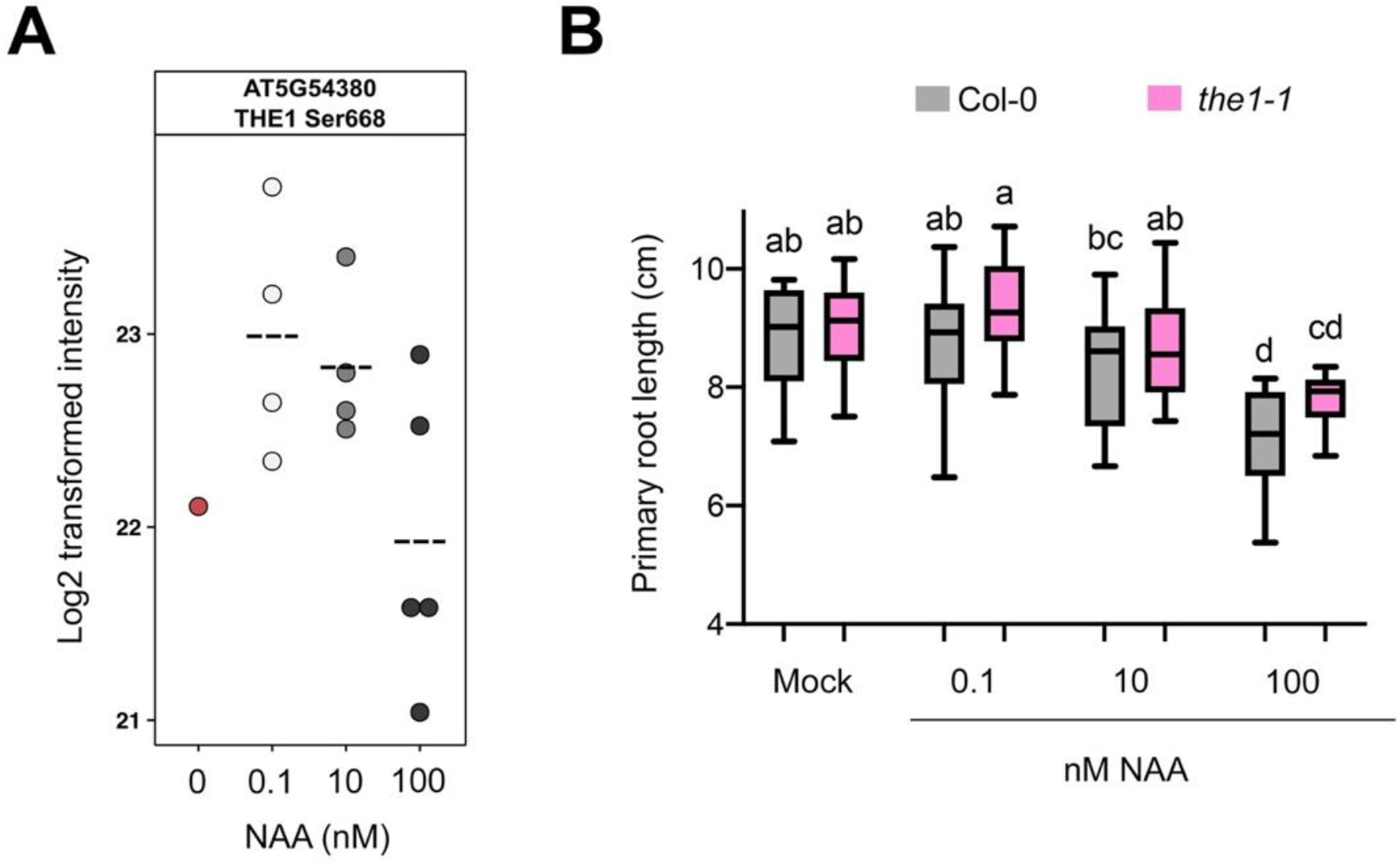
THE1 impacts auxin-controlled primary root growth. **(A)** THE1 phosphoprofile for the TGPSLDQT(0.007)HV<u>S</u>(0.959)T(0.034)AVK phosphopeptide upon NAA treatment. Dashed line indicates mean. Each dot is a biological replicate. **(B)** Primary root growth of *the1-1* at different NAA concentrations (n = 18-30 seedlings) at 11 days after germination. Boxplots show average with Tukey-based whiskers. Letters indicate significant difference according to two-way ANOVA with Tukey post-hoc test (p < 0.05).

Eukaryotic MAPK cascades act downstream of receptors or sensors to transduce various extracellular stimuli (Kim et al., 2012; Meng and Zhang, 2013; Singh and Jwa, 2013). Our list of differential phosphorylated proteins contained several members of MAPK cascades: two MAPKKK7, MKK2 and MAPK8 (**Supplementary Table S8**). In the *Arabidopsis* genome based on sequence homology there are approximately 60 MAPKKKs, 10 MAPKKs and 20 MAPKs and some of the members of this kinase family are known to be functionally redundant (Meng and Zhang, 2013). Given that complexity, we focused our attention on the “bottleneck” of MAPK cascades, a member of the MAPKKs that we identified in our study, namely MKK2. Previously, MKK2 has been associated with biotic and abiotic stresses in several studies (Teige et al., 2004; Kong et al., 2012; Wang et al., 2014; Gao et al., 2017). Our data showed that phosphorylation of MKK2 at the conserved Thr^31^ was identified only in samples treated with 0.1 and 100 nM of NAA (**Figure 4A and Supplementary Figure S12**). The conserved Thr^31^ in the MKK2 protein suggested that the phosphorylation of this residue may play an important regulatory role, for example in banana during cold stress response (Gao et al., 2017). Considering that MKK2 is closely related to and has high sequence similarity to MKK1 (Gao et al., 2008), we included both in our further analysis. We evaluated primary root growth of loss-of-function mutants *mkk1-2* (SALK_027645) and *mkk2-1* (SAIL_511_H01) in response to different auxin concentrations. At 9 days after germination *mkk2-1* demonstrated higher sensitivity to the growth promoting auxin concentration (**Figure 4B**). However, it should be noted that the difference in primary root length for *mkk2-1* is only 8.0 % for 0.1 nM NAA compared to Col-0. However, this might be explained by functional redundancy of these two members of MAPK cascade (Gao et al., 2008). Unfortunately, the *mkk1 mkk2* double mutant is severely impaired in its growth and development and often displays lethality at seedling stage (Gao et al., 2008; Qiu et al., 2008), making it difficult to examine primary root growth responses to different auxin concentrations. At the same time, treatment with a strongly growth inhibiting NAA concentration (100 nM) did not reveal any differences in *mkk1*-2 or *mkk2-1* primary root growth compared to wild type (**Figure 4B**).

**Figure 4.**
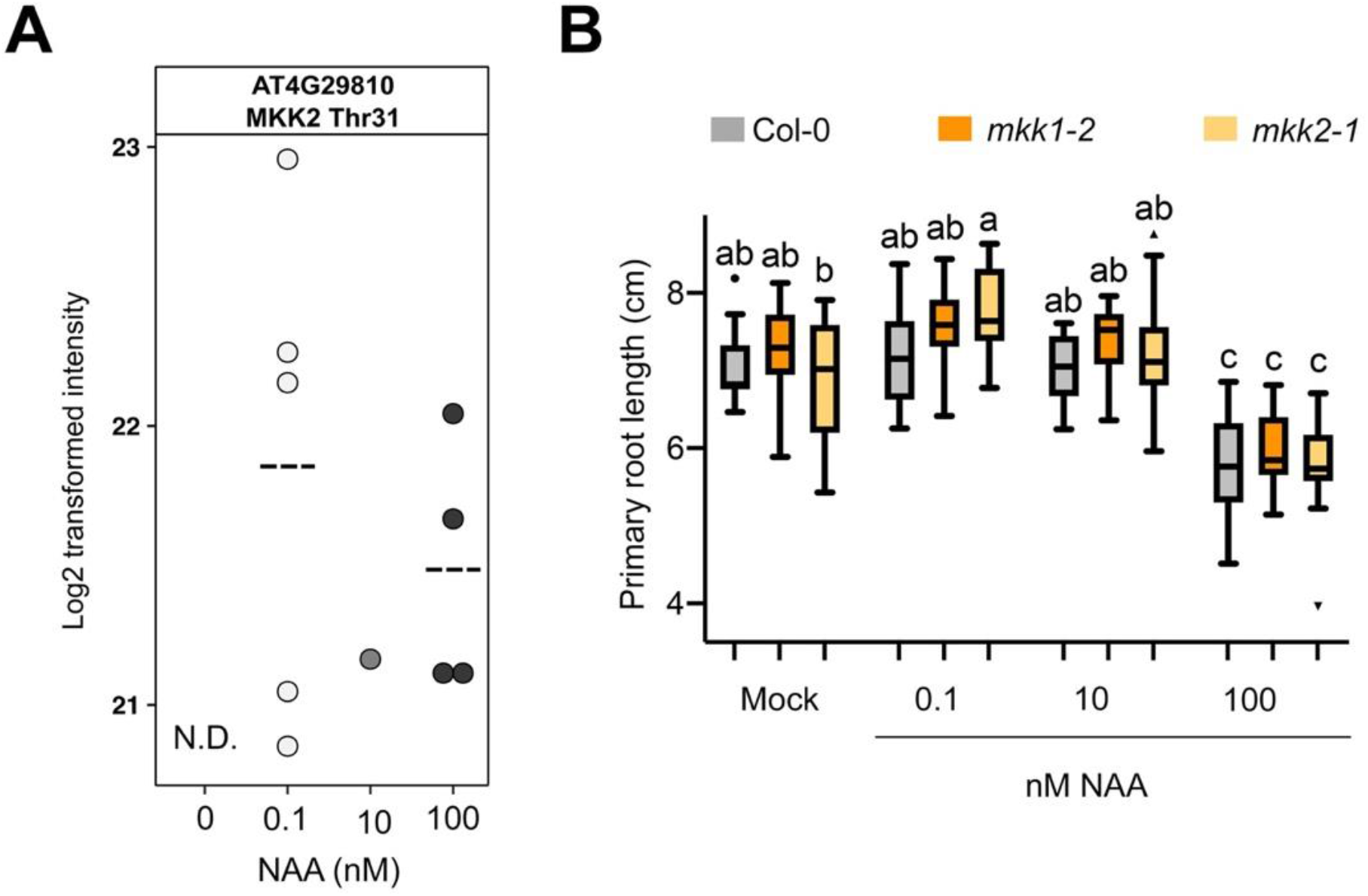
MKK2 impacts auxin-controlled primary root growth. **(A)** MKK2 phosphoprofile for the FLT(0.001)QS(0.031)GT(0.968)FK phosphopeptide upon NAA treatment. Dashed line indicates mean. Each dot is a biological replicate. N.D., not detected. **(B)** Primary root growth of *mkk1-2* and *mkk2-1* at different NAA concentrations (n = 10-16 seedlings) at 9 days after germination. Boxplots show average with Tukey-based whiskers and outliers. Letters indicate significant difference according to two-way ANOVA with Tukey post-hoc test (p < 0.05).

In addition, we observed an auxin concentration-dependent downregulation of PROHIBITIN3 (PHB3) protein levels (**Supplementary Figure S13A**). PHB3 coordinates cell division and differentiation in the root apical meristem through restricting the spatial expression of ETHYLENE RESPONSE FACTOR (ERF) transcription factors 115, 114, and 109 (Kong et al., 2018). Indeed, a *phb3* mutant showed a short primary root (**Supplementary Figure S13B**), indicating that selecting candidates based on a differential protein level allows identifying novel primary root growth regulators.

### The RALF34-THE1 module controls auxin-dependent primary root growth

RALF peptides have previously been shown to impact H^+^-ATPase (AHA) phosphorylation, which results in small changes in pH that affect root cell elongation (Haruta et al., 2014; Dressano et al., 2017). Indeed, we find extensive auxin concentration-dependent regulation of AHA1 and AHA2 phosphorylation, on residues that affect pump activity (Falhof et al., 2016) in our data set (**Supplementary Figure S9D-J**). In addition, it was shown that THE1 is a pH-dependent receptor for the RAPID ALKALINIZATION FACTOR 34 (RALF34) peptide and that this signalling module has a role in the fine-tuning of lateral root initiation and in regulating hypocotyl elongation (Gonneau et al., 2018). We first confirmed, using a transcriptional reporter (*pRALF34::n3xRFP*) (Murphy et al., 2016), the strong *RALF34* expression in the entire root meristem, with the strongest signal in the root cap and epidermis (**Figure 5A**). To explore the role of RALF34 in primary root growth, we analyzed two T-DNA insertion lines in *RALF34* (*ralf34-1* and *ralf34-2*, which were previously described (Murphy et al. 2016). Compared to their respective controls, Col-0 and L*er*, both *ralf34-1* (10.3 %) and *ralf34-2* (18 %) displayed a significantly longer primary root (**Figure 5B**). To establish the responsiveness of *ralf34-1* to auxin, we exogenously applied the synthetic phytohormone NAA to the mutant and examined their sensitivity. At 10 days post germination *ralf34-1* seedlings displayed slightly (but only significantly for 50 nM NAA; 13.2 %) longer primary roots than wild type at all NAA concentrations (**Figure 5C**), indicating a reduced sensitivity to growth-repressing auxin concentrations. In addition, we observed that primary root growth in the *ralf34-1* mutant after a 6-hour treatment with 10 nM IAA showed mildly decreased auxin sensitivity, in comparison to Col-0 (**Figure 5D**). This indicates that RALF34, possibly redundantly with other RALFs, is a component downstream of auxin exerting its effect on primary root growth.

**Figure 5.**
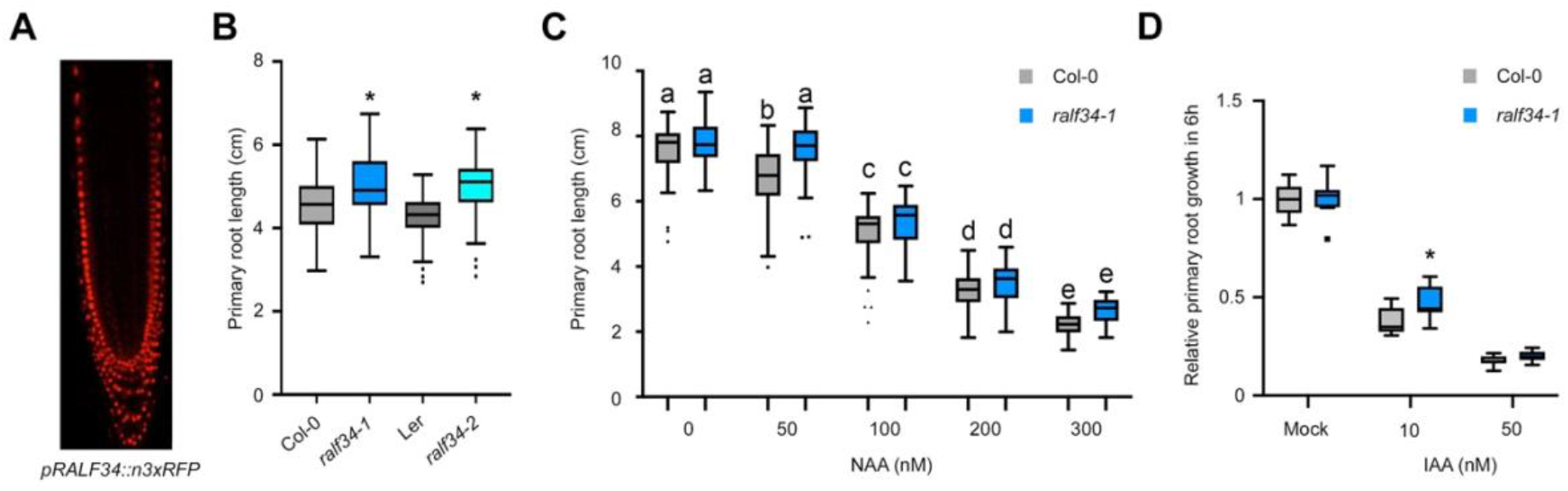
RALF34 impacts (auxin-controlled) primary root growth. **(A)** *RALF34* expression in the primary root tip as visualized through *pRALF34::n3xRFP*. **(B)** Primary root length (cm) of 7 days post germination seedlings: wild type Col-0 (n=133), L*er* (n=40), and RALF34 T-DNA insertion mutants *ralf34-1* (n=104) and *ralf34-2* (n=120). Boxplots with Tukey-based whiskers and outliers show data from 3 biological replicates. Asterisks indicate statistical significance (p < 0.001) based on Student’s t-test when T-DNA line is compared to its control. **(C)** Primary root length (cm) of seedlings 10 days post germination in absence (EtOH) or presence of various quantities of auxin (NAA) for Col-0 wild type (n= 63, 59, 61, 62, and 41, respectively) and *ralf34-1* (n= 48, 56, 41, 43, and 24, respectively). Boxplots with Tukey-based whiskers and outliers show data from 2 biological replicates. Letters indicate significant difference according to two-way ANOVA with Tukey post-hoc test (p < 0.05). **(D)** Normalized growth for 6 hours (to mock condition) for Col-0 and *ralf34-1* treated with indicated auxin concentration (IAA). Boxplots with Tukey-based whiskers and outliers (7 < n < 11). Asterisks indicate statistical significance (p < 0.05) based on Student’s t-test when *ralf34-1* is compared to Col-0.

### Proteome profiling identifies molecular changes downstream of MKK1 and MKK2

To our knowledge, MKK1 and MKK2 have not been previously implicated in primary root growth and auxin biology. Therefore, to gain insight in the molecular processes affected by a loss of function of MKK1 or MKK2, we performed proteome profiling on respective mutants in the absence of auxin. For this, total roots of 11-days-old seedlings of *mkk1-2* and *mkk2-1* mutants were harvested and proteome analysis was performed in 3 biological replicates. Statistical analysis revealed 172 differentially abundant proteins (**Figure 6 and Supplementary Table S9**). In addition, 12 proteins were identified as ‘unique’ (because these were not detected in at least one of the genotypes) (**Figure 6 and Supplementary Table S9**). Six proteins were absent in both *mkk1-2* and *mkk2-1* proteomes, namely peptidyl-prolyl cis-trans isomerase CYP18-1 (AT1G01940), WPP DOMAIN-INTERACTING PROTEIN 2 (WIT2, AT1G68910), L-GALACTONO-1,4-LACTONE DEHYDROGENASE (GLDH, AT3G47930), NASCENT POLYPEPTIDE-ASSOCIATED COMPLEX SUBUNIT ALPHA-LIKE PROTEIN 2 (NACA2, AT3G49470), Mitochondrial import receptor subunit TOM7-1 (TOM7-1, AT5G41685) and Ribulose bisphosphate carboxylase large chain (RBCL, ATCG00490) (**Supplementary Figure S14**). Two proteins were absent in the *mkk2-1* proteome: AUXIN RESISTANT 1 (AXR1, AT1G05180) and SERINE/THREONINE PROTEIN PHOSPHATASE 2A 55 kDa regulatory subunit B beta isoform (PP2AB2, AT1G17720 (**Supplementary Figure S14**).

**Figure 6.**
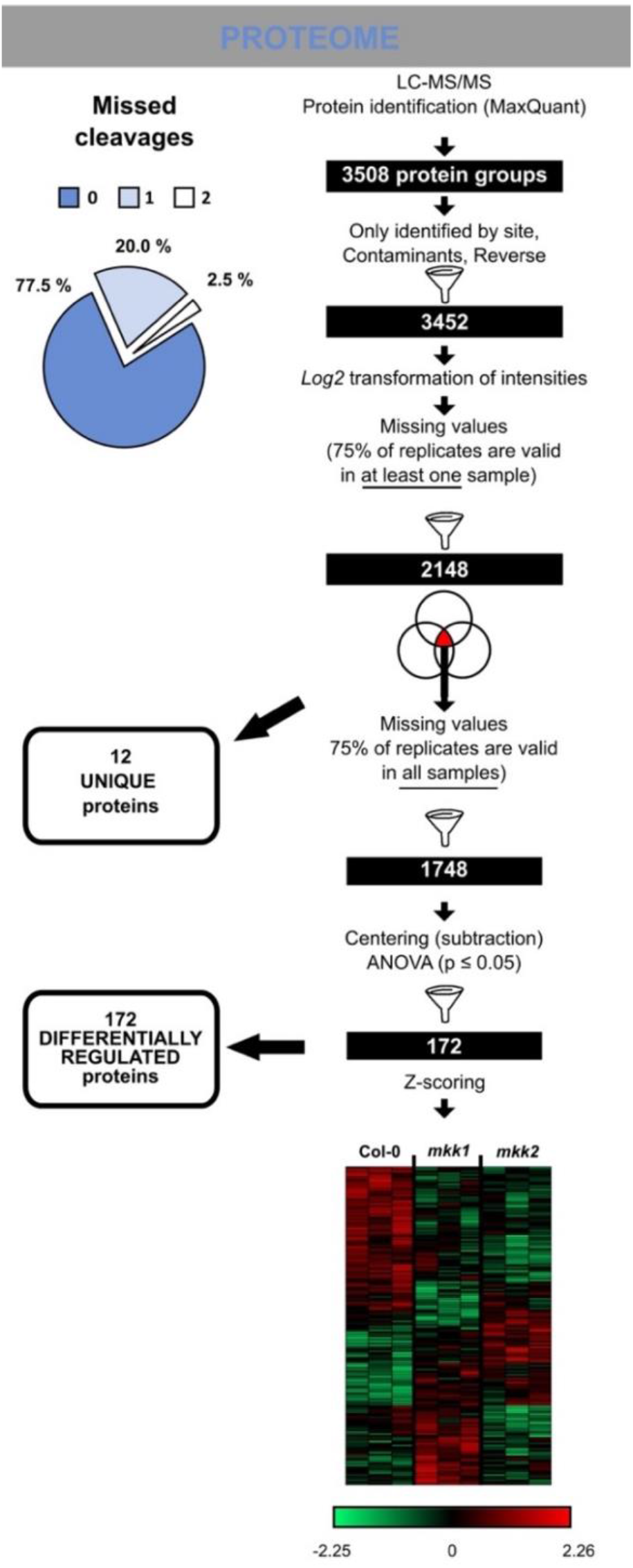
Protein changes in *mkk1-2* and *mkk2-1* mutants. Workflow illustrating the steps to obtain a reliable set of proteins or phospho-sites following LC-MS/MS. Venn diagrams indicate steps where ‘unique’ proteins/phosphosites (with corresponding numbers) were filtered out from the statistical analysis. Heatmap represents a hierarchical clustering of statistically significant proteins based on Pearson correlation. Centered Z-scored log2-transformed intensity values on heatmap are color-coded according to provided color gradient scale.

In *Arabidopsis* leaves the MKK2-MPK10 module regulates vein complexity by altering polar auxin transport efficiency (Stanko et al., 2014). In the same study, it was found that the *mpk10* mutant has an overlapping phenotype with some mutants in auxin-related genes, for example *AUXIN-RESISTANT 1 (AXR1)* (Stanko et al., 2014). Interestingly, our data revealed that AXR1 was one of the proteins absent in the root proteome of *mkk2-1* (**Figure 7A**). AXR1 mediates auxin response by activating the Skp-Cullin-F-box SCF E3 ubiquitin ligase complex that targets the AUX/IAA repressors of auxin response for ubiquitination and degradation (Dharmasiri et al., 2007). Lack of *AXR1* leads to resistance to growth inhibiting concentrations of auxin in *Arabidopsis* roots (Lincoln et al., 1990) and additionally to shorter root hairs that could indicate its role in cell elongation (Pitts et al., 1998). Indeed, analysis of *axr1-30* mutant revealed a significant increase of primary root length (**Figure 7B**).

**Figure 7.**
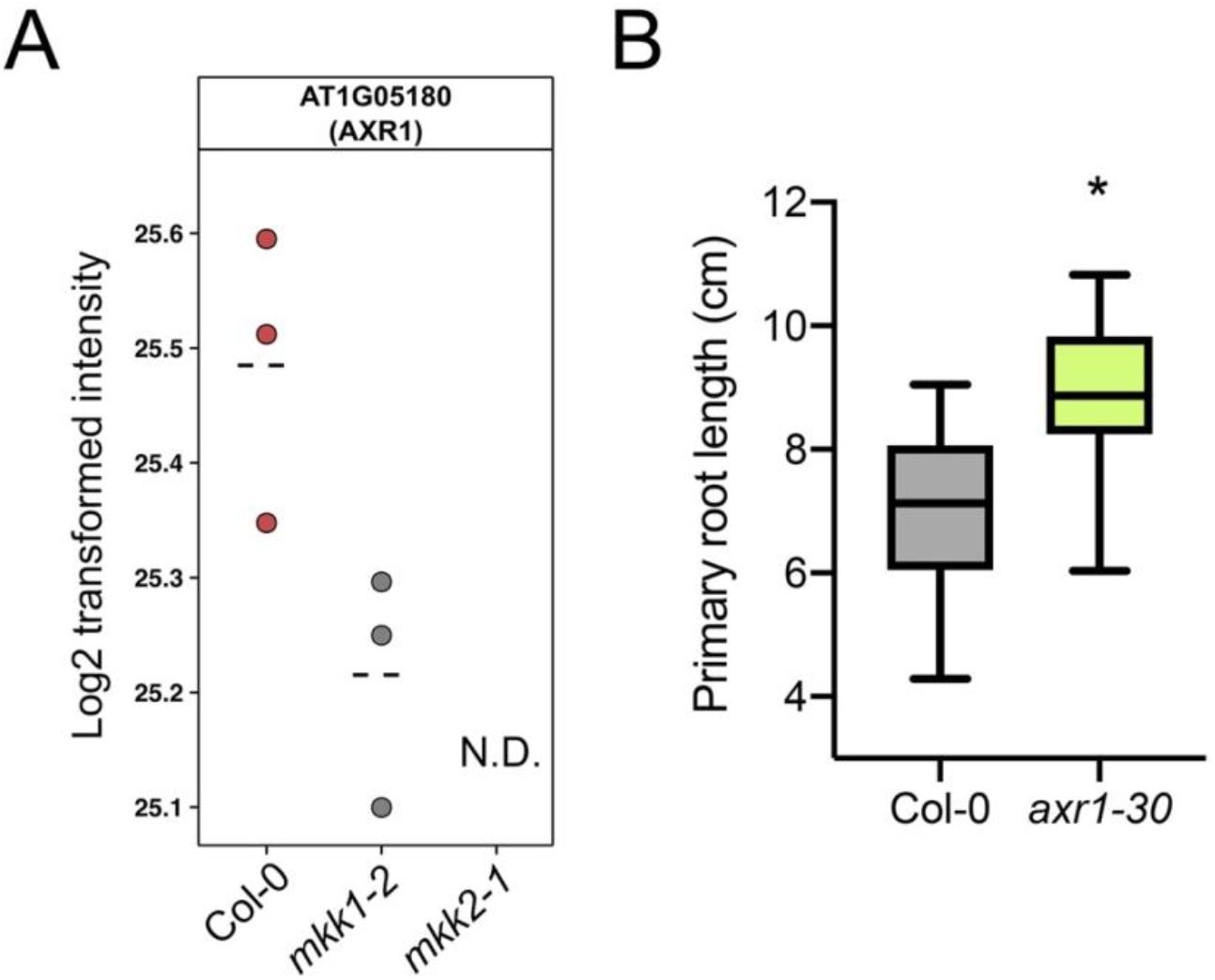
MKK1 and MKK2 affect AXR1 levels. **(A)** AXR1 protein profile in *mkk1-2* and *mkk2-1*. Dashed line indicates mean. Each dot is a biological replicate. N.D., not detected. **(B)** Primary root growth of *axr1-30* (n = 31-35 seedlings) at 11 days after germination. Boxplots show average with Tukey-based whiskers. A Student’s t-test revealed a significant difference (p < 0.05).

Among the differentially abundant proteins we identified, some proteins that are involved in auxin biosynthesis (**Supplementary Table S9)**. In both *mkk1-2* and *mkk2-1* mutants PHOSPHORIBOSYLANTHRANILATE TRANSFERASE 1 / TRYPTOPHAN BIOSYNTHESIS 1 (PAT1/TRP1, AT5G17990) was more than 3-fold downregulated. In addition, in both *mkk1-2* and *mkk2-1* mutants TRYPTOPHAN SYNTHASE ALPHA CHAIN / TRYPTOPHAN-REQUIRING 3 (TSA1/ TRP3, AT3G54640) was downregulated, 2.2- and 4.3-fold, respectively. TSA1 contributes to the tryptophan-independent indole biosynthesis, and possibly to auxin production (Radwanski et al., 1995; Radwanski et al., 1996; Rutherford et al., 1998; Ouyang et al., 2000). The *trp3-1* and *trp2-1* mutants, defective in tryptophan synthase α (TSA) and tryptophan synthase β (TSB) subunits, respectively, accumulate higher levels of IAA than the wild type despite containing lower Trp levels (Normanly et al., 1993; Ouyang et al., 2000). In addition, two nitrilases were identified that catalyse the terminal activation step in indole-acetic acid biosynthesis. NITRILASE 1 (NIT1, AT3G44310), a predominantly expressed nitrilase, was more than 2.5-fold downregulated in *mkk2-1*. NITRILASE 3 (NIT 3, AT3G44320), an enzyme that is able to convert indole-3-acetonitrile to indole-3-acetic acid (IAA) (Bartling et al., 1992; Bartling et al., 1994), was downregulated in both *mkk1-2* and *mkk2-1* mutants, 3.3- and 3.4-fold, respectively.

Although no large changes in phenotype and growth responses in response to auxin were observed for *mkk1-2* and *mkk2-1* (**Figure 4**), the root proteome data indicates altered auxin signalling and biosynthesis. Previously MKK2 was found to be involved in regulation of polar auxin transport efficiency in leaves (Stanko et al., 2014). However, our data suggested that in roots MKK2 and/or MKK1 seem to be also involved in signalling cascade regulating auxin biosynthesis. To further explore this, validation of proteome data, for example, analysis of auxin level and expression of genes involved in auxin biosynthesis in roots of *mkk1* and *mkk2* mutants is required.

## CONCLUSION

In this study, a MS-based phosphoproteomic approach was used to identify and characterise global auxin-mediated changes in protein abundance and phosphorylation underlying root growth promotion and inhibition. In roots, canonical auxin signalling leading to modulation of gene transcription and protein abundance is debatable with respect to rapid growth responses (Tan et al., 2007; Fendrych et al., 2016). Such rapid growth responses are more likely determined by plasma membrane depolarisation, Ca^2+^- and pH signalling and phosphorylation (Monshausen et al., 2011; Takahashi et al., 2012; Vanneste and Friml, 2013; Shih et al., 2015; Barbez et al., 2017). Our data also support a mechanism of non-transcriptional auxin-mediated growth regulation. Specifically, upon auxin treatment growth responses were much more pronounced and had more complex profiles at the level of protein phosphorylation rather than changes in protein abundance. In view of other cellular mechanisms downstream of auxin, such as microtubule re-arrangements (Chen et al., 2014) or vacuolar fragmentation (Löfke et al., 2015; Scheuring et al., 2016), we also detected several proteins related to these processes (**Supplementary Tables S1-S2**). From the phosphoproteome data we furthermore pinpointed some (novel) growth regulators from members of receptor-like kinases and MAP kinases. Our results together with previously published studies suggest that auxin, H^+^-ATPases, cell wall modifications and cell wall sensing receptor-like kinases are tightly embedded in a pathway regulating cell elongation (Hématy et al., 2007; Gonneau et al., 2018; Schoenaers et al., 2018). Furthermore, our study assigned a novel role to MKK2 in primary root growth and as a (potential) regulator of auxin biosynthesis and signalling, and supports the likely importance of MKK2 Thr^31^ phosphorylation site for growth regulation in the *Arabidopsis* root tip. Further studies will shed more light on the physiological significance of these proteins and the role the phosphorylation evens play.

## MATERIAL AND METHODS

### Plant materials and Growth conditions

The following lines were use: *mkk1-2* (SALK_027645) and *mkk2-1* (SAIL_511_H01) (Gao et al., 2008), *the1-1* (Hématy et al., 2007), *ralf34-1* and *ralf34-2* (Murphy et al. 2016). All T-DNA insertion lines used in current study were in Col-0 background apart from *ralf34-2* that was in L*er* background. Seeds were surface-sterilized and sown on half strength Murashige and Skoog (MS) medium, containing 1% (W/V) agarose and 0.8% sucrose, pH5.8. Seeds were stratified at 4°C in dark 2 days. After 2 days of stratification, seeds were germinated on vertically positioned Petri dishes in a growth chamber at 21°C under continuous light (100 µmol m^-2^ s^-1^ photosynthetically active radiation). *Arabidopsis* seedlings were grown vertically on half strength MS medium for indicated time after germination under above-mentioned conditions. For the *eru* mutant (SALK_083442C) seeds were surface-sterilized and sown on Gilroy medium (Wymer et al., 1997) with different concentrations of NAA for 6 days or on Gilroy medium without treatment for 4 days + 2 days on Gilroy medium with NAA after transfer. Gilroy medium always contained 0.8% phytagel and 1% sucrose at pH 5.7. Seeds were germinated on vertically positioned Petri dishes in a growth chamber with 16 h light/8 h dark conditions, at 22^°^C. For *phb3* (SALK_020707) (Van Aken et al., 2007), *A. thaliana* seeds were surface-sterilized by chlorine fumigation, and then held for two days at 4°C on solidified Murashige and Skoog (MS) medium before being transferred to a growth room providing a 16 h photoperiod and a constant temperature of 22°C.

### Primary root length analyses

Plates were scanned at 600 dpi resolution. Primary root length was measured using the NeuronJ plugin in Fiji package (https://fiji.sc). For *eru*, images were collected using a Nikon AZ100 multizoom macroscope for determination of root length using Image J. Data was analysed and visualized using R software environment (https://www.r-project.org; version 3.3.3).

### Scanner Growth Assay

4- or 5-day-old seedlings were transferred to the surface of solid ½ MS (+ 1% sucrose) 1% agar medium with IAA treatments as indicated in the 60 × 15 mm petri dishes. The sample with the petri dishes were fixed onto a vertically mounted flatbed scanner (Epson perfection V370) and seedlings were imaged through the layer of medium. Scan was taken automatically every hour using the AutoIt script described previously (Li et al., 2018) at 800 or 1200 dpi. The resulting image series were analysed using StackReg stabilization and the Manual Tracking plugin in ImageJ.

### Data analysis

Data shown in graphs are average values of multiple biological repeats (as indicated in the figure legends). Images were processed with Inkscape.

### qPCR

1 cm roots tips of Col-0 seedlings vertically grown in presence of different NAA concentrations were collected at 11 days after germination. RNA was extracted using the RNeasy Mini Kit (Qiagen) according to the protocol provided by the manufacturer and eluted in 40 µL of RNAse-free water. The RNA concentration was measured on a NanoDrop ND-1000, and cDNA was synthesized using iScript™ cDNA Synthesis Kit (BIO-RAD) according to the protocol provided by manufacturer and diluted 8 times. Samples for qPCR were prepared by the JANUS® Automated Workstation (PerkinElmer) and qPCR was performed on a LightCycler® 480 (Roche). All samples include 5 biological and 2 technical repeats. Primers used are listed in table X. Data analysis and visualization was performed in qbase+ (v. 3.0) and R (https://www.r-project.org/).

### (Phospho)proteome analysis

For the NAA experiment protein extraction, phosphopeptide enrichment, LC-MS/MS analysis (Nikonorova et al., 2018) was performed on 1 cm roots tips of Col-0 seedlings vertically grown in the presence of different NAA concentrations. Proteome analysis of the *mkk1-2* and *mkk2-1* mutants was performed according to (Vu et al., 2016). Data analysis was performed according to (Nikonorova et al., 2018). For total normalization, we normalized all the phosphosites on all the proteins before statistical analysis in Perseus. Then on the normalized data we performed statistical analysis (ANOVA with p < 0.05) and clustered the output.

### Sequence alignment

Protein sequences of CrRLK1 family members were aligned using a progressive alignment algorithm (Feng and Doolittle, 1987) in order to create multiple sequence alignments with CLC DNA Workbench 7 using the following settings: Gap open cost (10), Extension cost (1), End gap cost (as any other), Alignment (very accurate).

### Accession numbers

All MS proteomics data have been deposited to the ProteomeXchange Consortium via the PRIDE partner repository (Vizcaíno et al., 2016) with the data set identifier PXD021267 [*Note for reviewers: Username: - Password: c3GLbXJJ*]. All of the MS/MS spectra can be accessed in MS-Viewer (Baker and Chalkley, 2014) using the search key rjfbkn8oqd (auxin phosphoproteome), vddjji7zda (auxin proteome) and 7mbvmfizrz (*mkk1 mkk2* proteome).

## ACKNOWLEDGMENTS

We thank the Nottingham Stock Centre for seeds, Frank Van Breusegem for the *phb3* mutant, and Herman Höfte for the *the1* mutant.

